# The effect of head orientation on vestibular signal-based modulation of paraspinal muscle activity during walking

**DOI:** 10.1101/2024.04.29.591089

**Authors:** Yiyuan C. Li, Koen K. Lemaire, Sjoerd M. Bruijn, Simon Brumagne, Jaap H. van Dieën

**Affiliations:** Department of Human Movement Sciences, Faculty of Behavioral and Movement Sciences, Vrije Universiteit Amsterdam, Amsterdam Movement Sciences, Amsterdam, The Netherlands; Department of Rehabilitation Sciences, KU Leuven, Leuven, Belgium

**Keywords:** Trunk control, Vestibulospinal reflexes, Paraspinal muscle, Electrical Vestibular Stimulation, Head orientation

## Abstract

Vestibulospinal reflexes play a role in maintaining the upright posture of the trunk. Head orientation has been shown to modify the vestibulospinal reflexes during standing. This study investigated how vestibular signals affect paraspinal muscle activity during walking, and whether head orientation changes these effects. Sixteen participants were instructed to walk on a treadmill for 8 min at 78 steps/min and 2.8 km/h in four conditions defined by the presence of electrical vestibular stimulation (EVS) and by head orientation (facing forward and facing leftward), while bipolar electromyography (EMG) was recorded bilaterally from the paraspinal muscles from cervical to lumbar levels. In both head orientations, significant phasic EVS-EMG coherence (*p* < 0.01) in the paraspinal muscles was observed at ipsilateral and/or contralateral heel strikes. Compared to walking with the head forward, a significant decrease (*p* < 0.05) was found in EVS evoked responses (i.e., EVS-EMG coherence and gain) when participants walked with the leftward head orientation, with which EVS induced disturbance in the sagittal plane. This overall decrease may be explained by less need of feedback control for walking stabilization in the sagittal plane compared to in the frontal plane. The decrease in coherence was only significant at the left lower vertebral levels and at the right upper vertebral levels around left heel strikes (*p* < 0.05). Together, these findings confirm the contribution of the vestibular afferent signals to the control of paraspinal muscle activity during walking and indicate that this control is changed in response to different head orientations.

**Key Point Summary:**

- Vestibulospinal reflexes simultaneously contribute to stabilizing the centre of mass trajectory and to maintaining an upright posture of the trunk.
- Head orientation, which challenges stability via altered visual, vestibular and proprioceptive signals, modifies vestibulospinal reflexes during standing.
- To explore the impact of head orientation during walking, we recorded bilateral surface EMG of cervical to lumbar paraspinal muscles, and characterized coherence, gain and delay between EMG and electrical vestibular stimulation, during walking with head facing forward and leftward.
- When walking with head facing leftward, vestibular stimulation caused disturbance in sagittal plane. Phasic response in paraspinal muscles with a significant smaller magnitude was found compared to facing forward.
- Our results agree with the idea that less feedback control is required for walking stabilization in the sagittal plane and confirm that vestibular afference modulates paraspinal muscle activity for trunk control during walking, and this modulation is changed in response to head orientation.

## Introduction

Controlling the orientation of the trunk is imperative to prevent falling when walking. With its large mass and relatively high location above the base of support, trunk has a large potential energy. Consequently, a small deviation from its upright orientation can produce a substantial destabilizing moment. To maintain the upright posture relies on feedback control, based on integrated visual, vestibular, proprioceptive and tactile inputs (Peterka, 2002 ; Goodworth & Peterka, 2009; Maaswinkel et al., 2014; Andreopoulou et al., 2015; Cofre Lizama et al., 2016; van Drunen et al., 2016). Disturbances of the vestibular system through electrical vestibular stimulation (EVS) have been widely used to investigate the contribution of vestibular signals to postural control, (Britton et al., 1993; Ali et al., 2003; Iles et al., 2007; Dakin et al., 2010; Forbes et al., 2013; Magnani et al., 2021). This stimulation, a small electrical current, activates the afferent nerve fibres of the vestibular organs and produces an illusion of self-movement, as a consequence of which a body sway opposite to the illusory movement is induced (Lund & Broberg, 1983; Mian & Day, 2009).

Coherence is a frequency-domain measure of linear association between signals. It varies between 0, indicating no association, and 1, indicating perfect association. EVS-EMG coherence thus indicates how strong muscle activity is coupled to afferent signals from the vestibular system. If the coupling is strong (i.e., coherence is significant), the frequency response function between the EVS signal and EMG activity can be estimated, which provides information on the gain and delay of the feedback from the vestibular system to muscle activity. Previously, we studied the coherence between EVS and paraspinal muscle activity from the C7 to L4 vertebral level while walking with the head facing forward. We showed that vestibular signals modulated paraspinal muscle activity (Li *et al*., 2024). Peak coherence was found at ipsilateral heel strike at lower vertebral levels, whereas it occurred at the contralateral heel strike at higher vertebral levels. These couplings appeared to be functional in maintaining an upright trunk orientation in the frontal plane.

In standing, EVS induced responses have been demonstrated to be craniocentric (Lund & Broberg, 1983; Mian & Day, 2014). For example, with the anode placed on the right mastoid, a rightward body sway will be induced by EVS when facing forward, while a backward sway will be evoked when the head is turned 90 degrees to the right (Lund & Broberg, 1983; Ali *et al*., 2003). A study by Forbes et al. reported that EVS-EMG coherence in the soleus muscle was maximal with the head facing 90 degrees to the left, while it diminished progressively when participants rotated their head towards the right, and finally decreased to zero when facing forward (Forbes *et al*., 2016). Thus, the direction of EVS induced disturbances can be manipulated by changing head orientation.

Head orientation also changes the effect of EVS on paraspinal muscle activity in standing. When facing sideways, EVS induced disturbances in the sagittal plane, with synergistic activity of the bilateral paraspinal muscles. Facing forward, EVS induced disturbances in the frontal plane, with bilateral paraspinal muscles acting as antagonists (Ali *et al*., 2003). To our knowledge, it is unknown if, and how, paraspinal muscles responses to EVS change as a consequence of head orientation during walking.

In this study, we investigated whether- and how vestibular signals influence paraspinal muscle activity during walking with the head facing forward and sideways. We expected that when facing sideways, illusion of self-movement would be induced mostly in the sagittal plane, causing the left and right paraspinal muscles to respond synergistically. Consequently, we hypothesized that EVS-EMG coherence would show a more symmetric pattern between the left and right side of the trunk and among the upper to lower vertebral levels, compared to facing forward. Since walking can be passively stable in the sagittal plane (Bauby & Kuo, 2000; O’Connor & Kuo, 2009), less feedback control may be required in the sagittal plane compared to the frontal plane. Hence, we also hypothesized that the EVS-EMG coherence and gain would show an overall decrease when facing sideways compared to facing forward. No impact of head orientation on the delay of the feedback from the vestibular system to muscle activity was expected.

## Methods

Part of the data presented in this study were reported earlier by Li et al (Li et al., 2024). Here, we briefly summarize the procedures.

### Participants

Sixteen healthy participants (7 male, 9 female) were recruited (Age: 23.5 ± 3.4 years, height: 1.71 ± 0.07 m, weight: 64.5 ± 13.2 kg). Participants were recruited only if they did not have any diagnosed orthopaedic or neurological disorders and did not use medication that can cause dizziness. Participants were instructed not to engage in intense physical exercise and to refrain from alcohol consumption 24 hours prior to the experiment. Two participants were excluded due to technical problems in the signal synchronization. Participants provided written informed consent.

The experimental design and procedures were approved by the VU Amsterdam Research Ethics Committee (VCWE-2022-126) and conformed to the Declaration of Helsinki, except for pre-registration.

### Electrical Vestibular Stimulation

Participants were exposed to a zero-mean stochastic EVS signal with a bandwidth from 0 to 25 Hz, peak amplitude of ± 5.0 mA, and duration of 8 minutes (Dakin *et al*., 2010) This signal was applied as an analogue signal through a digital-to-analogue converter (National Instruments Corp., Austin, USA) to an isolated constant-current stimulator (BIOPAC System Inc., Goleta, USA), which was connected to carbon rubber electrodes (9 cm^2^). The electrodes were coated with electrode gel (SonoGel, Bad Camberg, Germany) and were placed over the mastoid processes behind the ears.

### Ground Reaction Forces and Kinematics

Ground reaction forces (GRF) were measured at 1000 samples/s by force plates embedded within the split-belt treadmill (ForceLink b.v., Culemborg, the Netherlands). Kinematic data were collected using a 3D motion capture system (Northern Digital Inc, Waterloo Ont, Canada) at a sampling rate of 50 samples/s. Cluster markers were attached to the feet, shanks, thighs, pelvis, trunk, head, upper arms and forearms. Corresponding anatomical landmarks were digitized using a six-markers probe.

### Electromyography

A 64-channel REFA amplifier (TMSi, Oldenzaal, The Netherlands, CMRR: > 90 dB, Input impedance: > 100 MΩ, Gain: 26.55) was used to record the bipolar surface electromyography (EMG) at 2048 samples/s. After shaving, abrading, and cleaning the skin with alcohol, paired disposable self-adhesive Ag/AgCl surface electrodes (Ambu blue sensor, Ballerup, Denmark) were placed 2 to 3 cm lateral to the spinous processes at eight vertebral levels bilaterally, including the seventh cervical vertebra (C7), the third, fifth, seventh, nineth and twelfth thoracic vertebra (T3, T5, T7, T9, T12) and the second and fourth lumbar vertebra (L2, L4). A reference electrode was placed over the acromion process.

### Protocol

Before the experiment, participants were exposed to vestibular stimulation in a three-minutes-walk at 2.8 km/h and were instructed to align their steps to the beat of a metronome at 78 steps/min for familiarization. During the experiment, participants walked on the dual-belt treadmill at 2.8 km/h with a cadence of 78 steps/min for 8 minutes in six conditions. The conditions were defined by the presence of EVS (EVS and no-EVS), by head orientation: head forward (HF) or turned to the maximal comfortable position to the left (HL) and by step width (preferred step width: PSW and narrow step width: NSW). The order of the conditions was randomized for each participant. Participants received repeated verbal instruction to maintain their cadence and head orientation during the trials. Data from narrow step width conditions were not used in this study.

### Signal Analysis

The global coordinate system was defined as positive x laterally to the right, positive y-axis forward, positive z-axis vertically upward. To measure if, and how well participants performed the task of controlling head orientation and cadence, the time series of the head orientation was first calculated using Euler decomposition (z-y-x sequence) in the global coordinate system, then averaged over gait cycles at each normalized time point for each participant. To remove any offset in the local joint coordinate systems, orientations in the head leftward (HL) condition were corrected by subtracting the averaged orientation in walking with head forward without EVS for each participant. In the rest of this paper, we define the head orientation as 0 degree when head facing forward, positive angles as facing leftward, and negative angles as facing rightward.

The electromyography, force plate and vestibular stimulation signals were synchronized based on trigger signals. Gait events (i.e. heel-strikes and toe-offs) were calculated based on the centre of pressure (Roerdink *et al*., 2008). Cadence was calculated from the time between subsequent heel strikes of the same leg. EMG signals (sampled at 2048 Hz) were resampled to 1000 Hz to align with the sampling rate of the force plate and EVS signals. To remove EVS artifacts, a sixth order Butterworth high pass filter with a cut-off frequency of 100 Hz was applied on bipolar EMG signals, followed by rectification of the filtered EMG signals (Blouin *et al*., 2011; Forbes *et al*., 2014; Li *et al*., 2024). For each vertebral level and participant, rectified EMG was normalized to the maximum amplitude of the EMG signal of the averaged gait cycle during walking with preferred step width, while facing forward and without EVS. Then, based on right heel strikes, the EVS and normalized EMG signals were sliced into gait cycles. To avoid distortion in the coherence and gain calculation, each gait cycle was padded with data from the previous (50%) and subsequent (50%) cycle. The coherence and frequency response function (FRF) between EVS and EMG were calculated based on the continuous Morlet wavelet decomposition as described in Li et al (2024). The gain (1/milliampere, mA^-1^) and phase estimates (radian, rad) between EVS and EMG were defined as the modulus and the angle of frequency response function, respectively. Phase estimates were transformed into time lags (millisecond, ms) by dividing by their corresponding frequencies. The gain and delay were defined as the median of the gains and time lags across the frequencies and gait phases at peak coherence for each vertebral level and each participant.

### Statistical Analysis

To test whether the instructions were followed, repeated-measures ANOVAs with two factors: Head Orientation (HO: Head Forward (HF) and Head Left (HL)) × the Presence of EVS (EVS and No EVS) were performed on head orientation (yaw) and cadence. To determine the differences in average muscle activity between conditions, one-dimensional Statistical Parametric Mapping based repeated-measures ANOVAs were performed with two factors: Head Orientation × Presence of EVS (Pataky et al., 2016). To identify any substantial effects on muscle activity, significance was set as *p* value < 0.01.

Because coherence is naturally bounded between 0 and 1, the modified Fisher-Z transform was applied on coherence values before statistical analysis. For each participant, coherence exceeding 0.015, (i.e., p < 0.01), was defined as significant (Zhan *et al*., 2006). To identify whether the coupling between EVS and EMG significantly differed between head orientations, cluster-based permutation tests, using t-statistics and 2000 permutations, were applied to the coherence (Maris *et al*., 2007). The effects of Head orientation, Vertebral Level, and Side (L: Left and R: Right side of the trunk), and their interactions on delays and gains were tested with repeated-measures ANOVAs. Sphericity for repeated-measures ANOVAs was verified by Mauchly’s Test (*p* > 0.05). Greenhouse-Geisser corrections were applied when the assumption of sphericity was rejected. A Bonferroni correction was applied for the post hoc tests. A *p* value less than 0.05 was defined as significant. Statistical analyses were performed in MATLAB (2019a, The MathWorks, Natick, US).

## Results

Averaged over time, participants maintained a head orientation of 50.948 (± 6.088) degrees during walking facing leftward (HL) (Figure 1A). No significant effect of EVS on averaged head orientation was found (*p*= 0.193).

**Figure 1.**
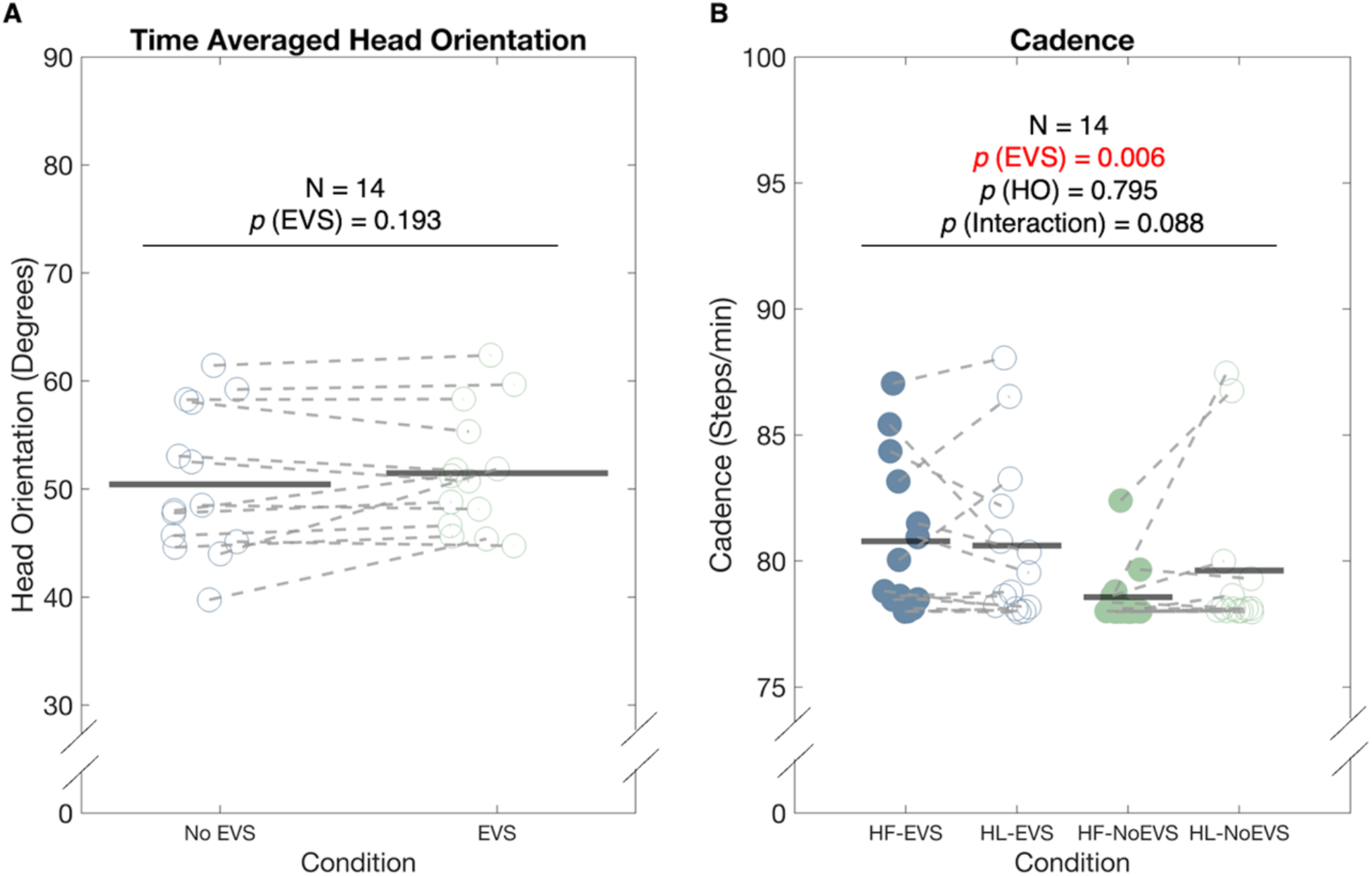
A: Time-averaged head orientation (in degrees) during walking with EVS (empty green dots) and without EVS (empty blue dots). B: Time-averaged cadence (in steps per minute) during walking in four conditions: Head Forward with EVS (HF-EVS, filled blue dots), Head Leftward with EVS (HL-EVS, empty blue dots), Head Forward without EVS (HF-No EVS, filed green dots) and Head Leftward without EVS (HL-No EVS, empty green dots). Each dot represents an individual data point. Black horizontal lines represent the averages across participants.

When walking without EVS and facing forward, participants maintained their cadence as instructed (78 steps/min vs 78.568 ± 1.159 steps/min). However, cadence significantly increased with the presence of EVS to 80.784 ± 2.953 steps/min (F (0.763,9.922) = 10.622, *p*= 0.006). No significant effects of Head Orientation or its interaction with EVS were found on cadence (Figure 1B).

Similar phasic activity of the paraspinal muscles was found in walking when facing leftward and forward (Figure 2). A significant effect of Head Orientation was only found for muscle activity at the right C7 level around heel strikes (Table 1), likely for maintaining the leftward head orientation during walking. Muscle activity at lower vertebral levels significantly increased with the presence of EVS. No significant effects of the interaction between Head Orientation and the presence of EVS on muscle activity were found (Table 1). Given these findings, it is reasonable to compare EVS-EMG coherence and gain between the two head orientation conditions below.

**Table 1.**
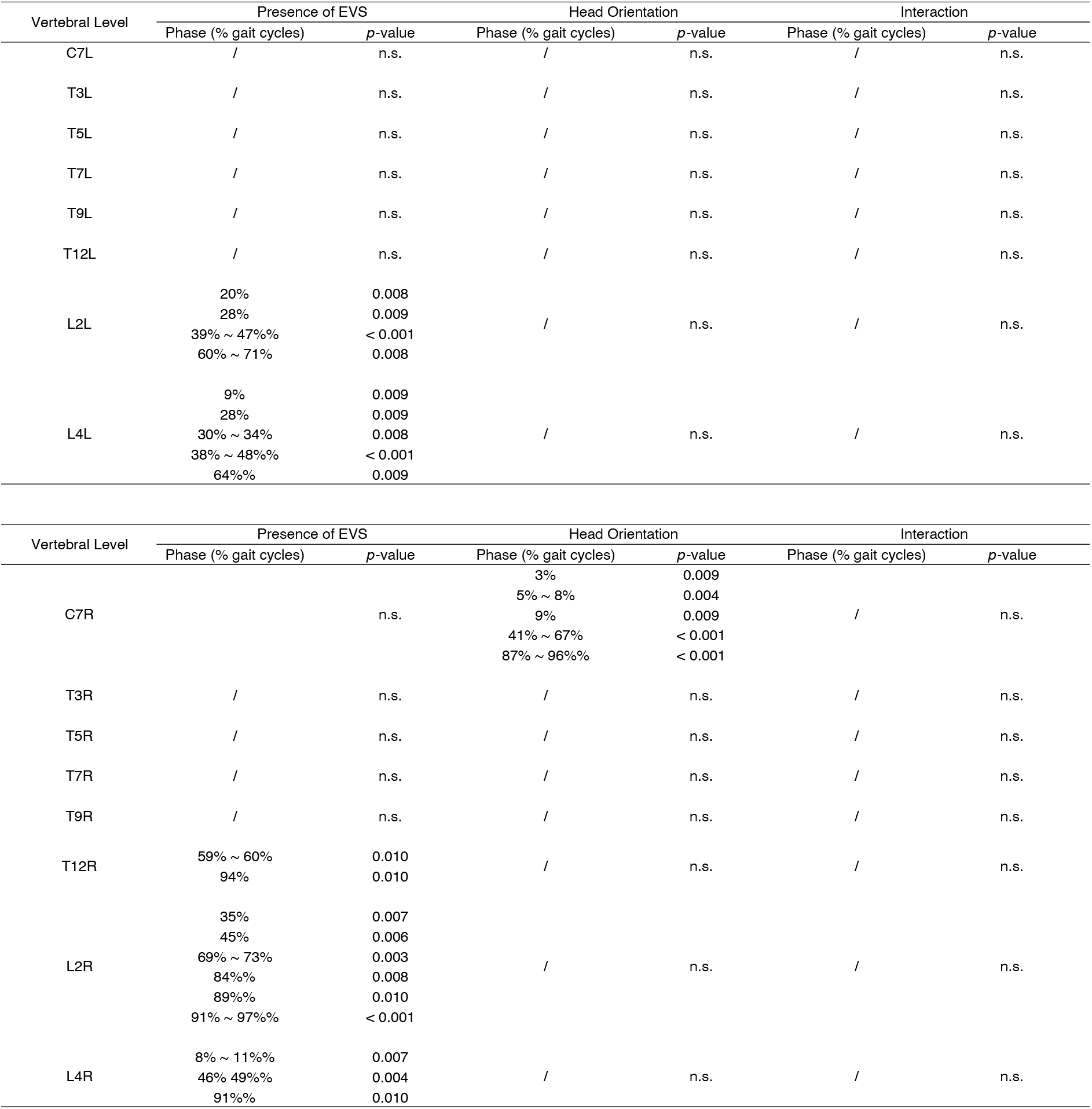
Summary of the effect of the presence of EVS and Head Orientation on Muscle Activity at Eight Vertebral Levels on the Left and Right Side Relative to the Trunk. Results from the One-dimensional Statistical Parametric Mapping based repeated-measures two-ways ANOVAs. The p-values indicate the probability that a suprathreshold cluster of the same spatiotemporal extent could have resulted from a random field process of the same smoothness as the observed residuals.

**Figure 2.**
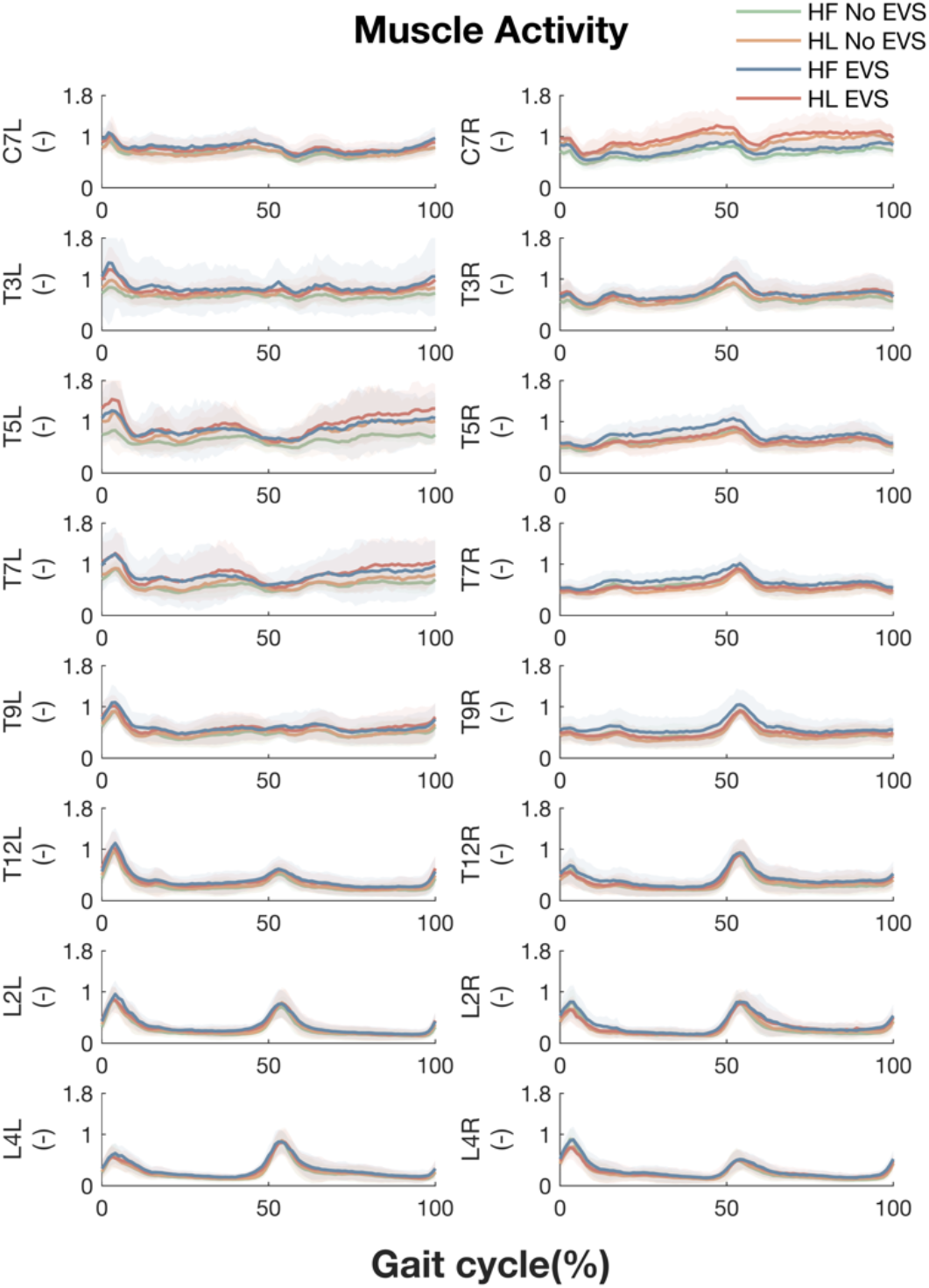
Averaged EMG linear envelopes of paraspinal muscles at all vertebral levels during walking with Head Forward without EVS (Green), with Head Leftward without EVS (Orange), with Head Forward with EVS (Blue) and with Head Leftward with EVS (Red). The left and right columns depict the left and right sides relative to the midline of the trunk. The X-axis is the time (normalized to gait cycle, starting on the right heel strike). The Y-axis indicates the amplitude of the normalized EMG. The lines and shadowed areas represent the means and standard deviations across participants, respectively.

### Phasic EVS-EMG Coupling in Walking with Head Leftward

The spatiotemporal pattern of EVS-EMG coherence was similar between walking with head forward (Li *et al*., 2024) and leftward. Significant EVS-EMG coherence was observed at all recorded sites and peaked around ipsilateral heel strike at lower vertebral levels and at contralateral heel strike at higher vertebral levels (Figure 3).

**Figure 3.**
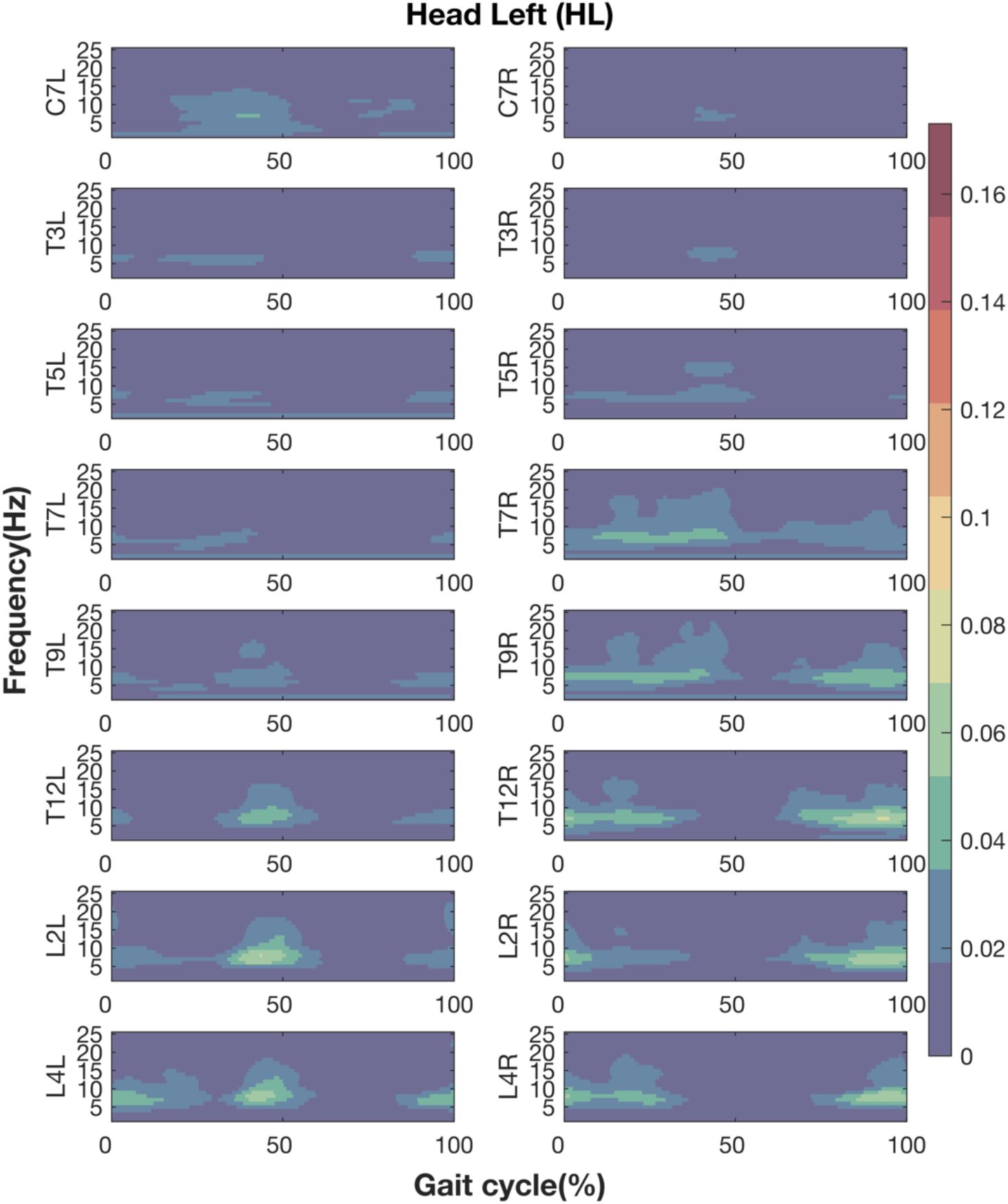
Coherence between EVS and EMG during walking with Head Leftward. Figures from top to bottom rows present the results at eight vertebral levels. The left and right columns depict the left and right sides relative to the midline of the trunk. The X-axis is the time (normalized to gait cycle, starting at the right heel strike). The Y-axis is the frequency ranging from 1Hz to 25 Hz. The magnitudes of coherence are indicated by the colour bar.

### Decrease in EVS-EMG Coupling Induced by Head Orientation

When walking facing leftward, EVS-EMG coupling significantly decreased around left heel strike at vertebral levels from T9 to L4 on the left side of the trunk and at vertebral levels from T3 to T5 level on the right side of the trunk (Figure 4).

**Figure 4.**
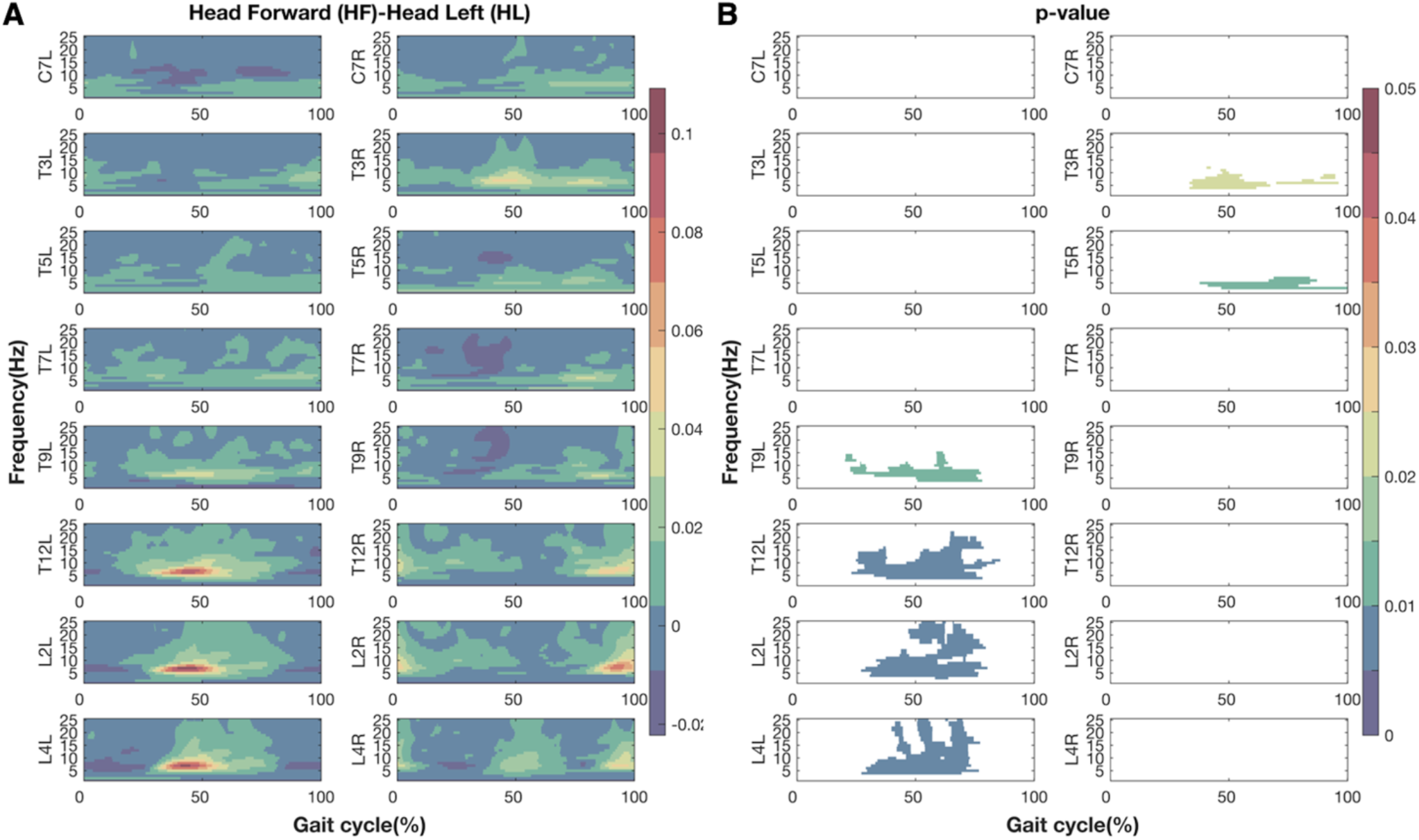
The differences in coherence (A) and associated p-values (B) between two head orientations (Head Forward-Head Leftward) during walking with EVS. Figures from top to bottom rows present the results at eight vertebral levels. The left or right columns represents the left or right sides relative to the mid-sagittal line of trunk. The X-axis is the time (normalized to gait cycle, starting on the right heel strike). The Y-axis is the frequency ranging from 1Hz to 25 Hz. The magnitudes are indicated by the colour bar. For clarity, only p-values lower than 0.05 are presented.

### Asymmetric EVS-EMG Coupling between the Left and Right Side of the Trunk

To further explore the asymmetric decrease in EVS-EMG coherence during walking with the head turned to the left, Statistical Parametric Mapping based paired t-tests were performed on the EVS-EMG coherence between the left and right side of the trunk, aligned at ipsilateral heel strikes (Figure 5A). During walking with the head turned left, EVS-EMG coherence was significantly higher on the left at C7 in early ipsilateral stance and late contralateral stance, and at L4 around contralateral heel strike. However, coherence was significantly lower at T9 and T12 during ipsilateral stance, and during contralateral stances at T12 (Figure 5B). Taken together, these effects confirm the asymmetric effects of head rotation on EVS-EMG coupling.

**Figure 5.**
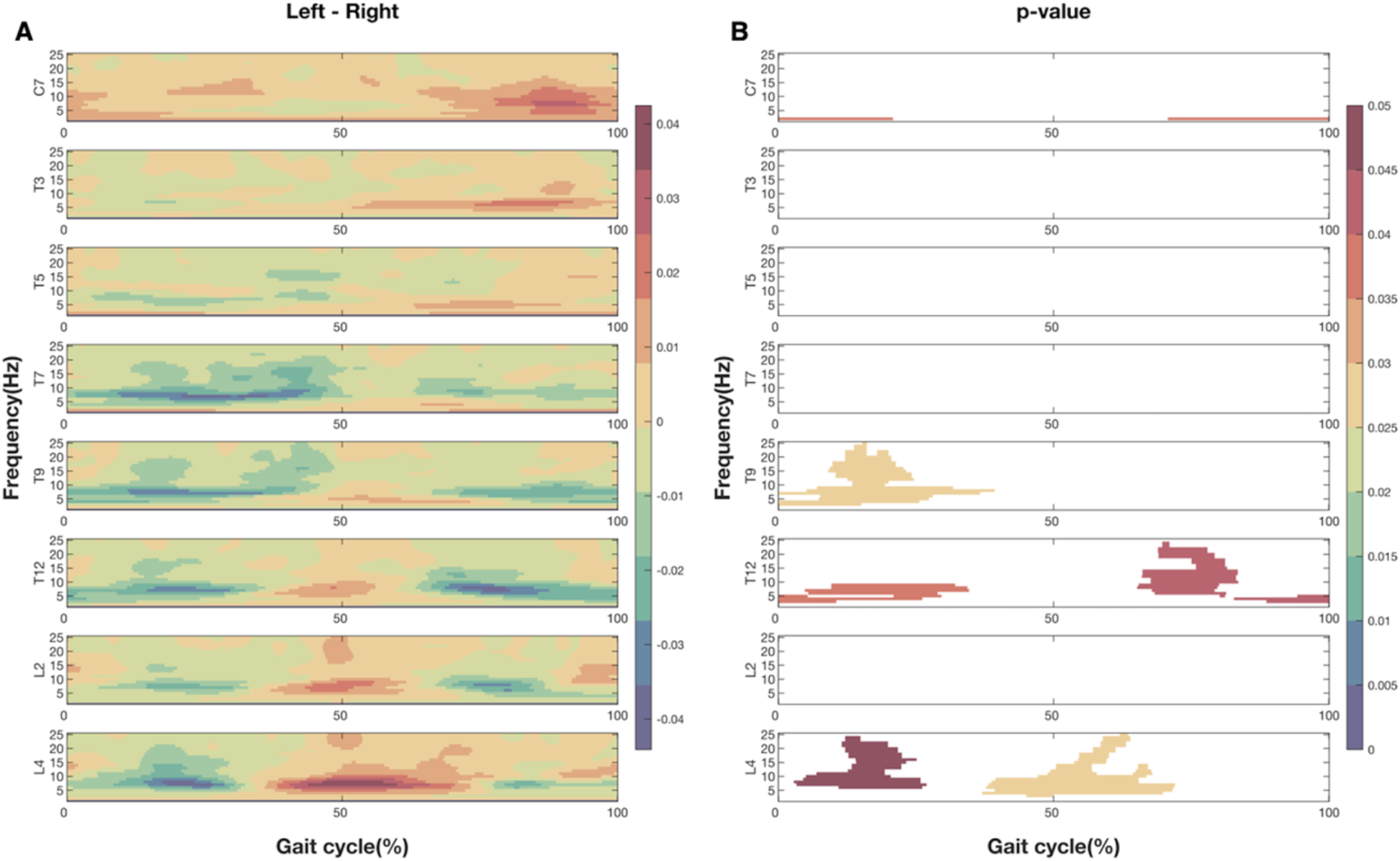
The differences in coherence (A) and associated p-values (B) between EVS and EMG during walking with head leftward between the left and right side of trunk (Left - Right). Figures from top to bottom rows present the results at eight vertebral levels. The X-axis is the time (normalized to gait cycle, starting at ipsilateral heel strike). The Y-axis is the frequency ranging from 1Hz to 25 Hz. The magnitudes are indicated by the colour bar. For clarity, only p-values lower than 0.05 are presented.

### Leftward Head Orientation Induced Decrease in Gain

Gain was significantly higher when walking facing forward than facing leftward (F (0.079, 1.027) = 12.883, *p*= 0.003) (Figure 6A & 6B, Table 2). No significant effect of either Vertebral Level, Side nor of any of the interactions was found on gain.

**Table 2.** Summary of the three-way ANOVA test (Head Orientation × Vertebral Levels × Side)

| Gain |  |  |  |
| --- | --- | --- | --- |
|  | N | F | p |
| Vertebral Levels | 14 | $F(0.553, 7.189) = 1.056$ | $p = 0.372$ |
| Head Orientations | 14 | $F(0.079, 1.027) = 12.883$ | $p = 0.003$ |
| Sides | 14 | $F(0.079, 1.027) = 1.483$ | $p = 0.245$ |
| Vertebral Levels × Head Orientations | 14 | $F(0.553, 7.189) = 1.840$ | $p = 0.190$ |
| Vertebral Levels × Sides | 14 | $F(0.553, 7.189) = 1.035$ | $p = 0.386$ |
| Head Orientations × Sides | 14 | $F(0.079, 1.027) = 0.018$ | $p = 0.895$ |
| Vertebral Levels × Head Orientations × Sides | 14 | $F(0.553, 7.189) = 1.598$ | $p = 0.213$ |
| Phase |  |  |  |
|  | N | F | p |
| Vertebral Levels | 14 | $F(1.379, 17.927) = 2.194$ | $p = 0.087$ |
| Head Orientations | 14 | $F(0.197, 2.561) = 0.161$ | $p = 0.695$ |
| Sides | 14 | $F(0.197, 2.561) = 0.091$ | $p = 0.768$ |
| Vertebral Levels × Head Orientations | 14 | $F(1.379, 17.927) = 0.423$ | $p = 0.753$ |
| Vertebral Levels × Sides | 14 | $F(1.379, 17.927) = 1.409$ | $p = 0.257$ |
| Head Orientations × Sides | 14 | $F(0.197, 2.561) = 0.345$ | $p = 0.567$ |
| Vertebral Levels × Head Orientations × Sides | 14 | $F(1.379, 17.927) = 0.717$ | $p = 0.559$ |

**Figure 6.**
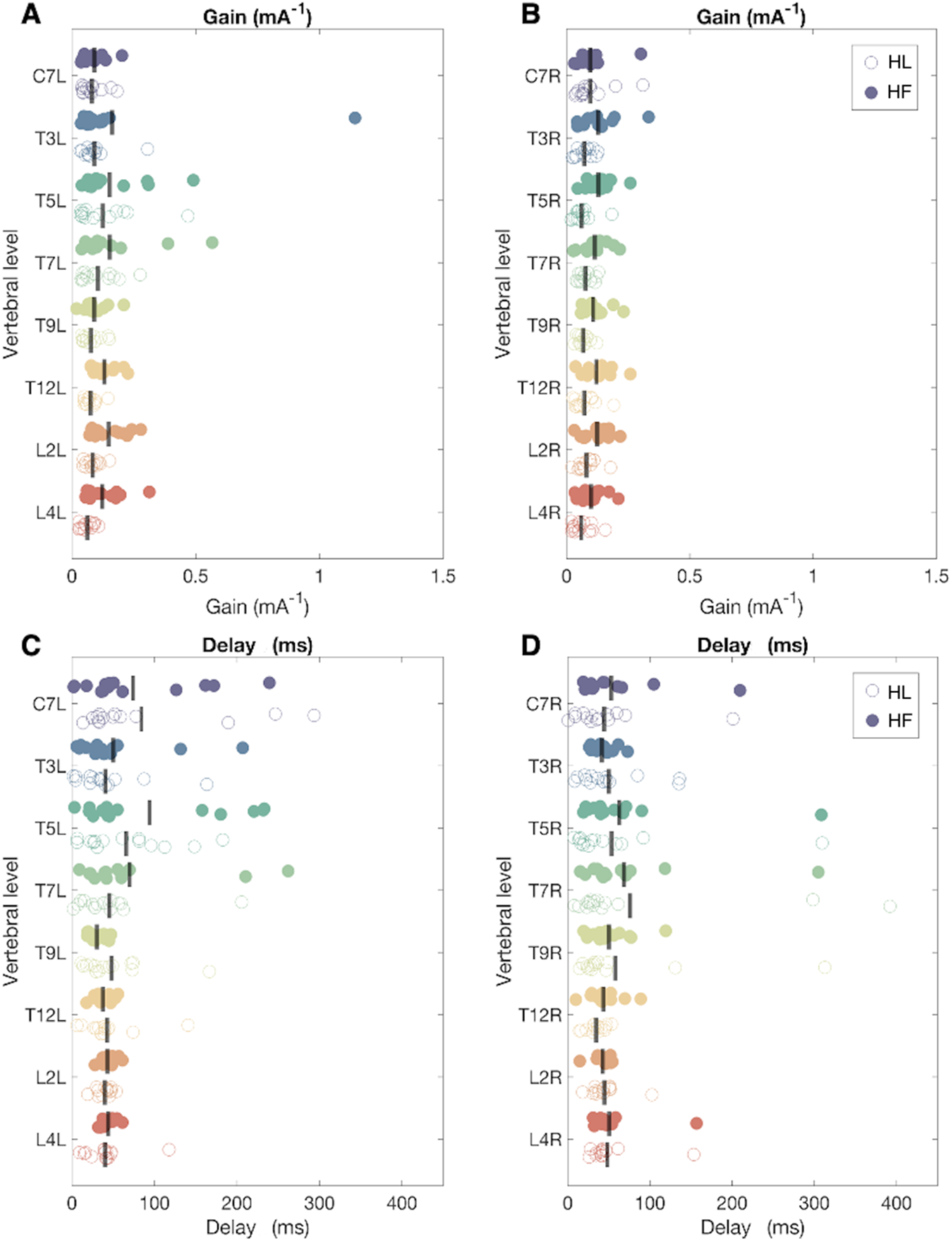
Gain and Delay Estimates between EVS and EMG. The Y-axis represents vertebral levels (C7, T3, T5, T7, T9, T12, L2 and L4) on the left (A & C) and right side (B & D) of the trunk. In panel A and B, the X-axis represents the magnitude of the gain (mA^-1^). In Panel C and D, the X-axis represents the delay (ms) between EVS and EMG. Each filled dot indicates the individual data during walking with head forward (HF). Each empty dot indicates the individual data during walking with head leftward (HL).

### No Effect of Head Orientation on Delays

As expected, no significant effects on delays of Head Orientation, Vertebral Level, Side or any of the interactions were found (Figure 6C & 6D, Table 2).

## Discussion

Using stochastic EVS, we compared how vestibular signals influence paraspinal muscle activity, from cervical to lumbar level, while walking with the head facing forward and sideways, respectively. Significant phasic coherence between the EVS and paraspinal muscle activity was observed when walking with the head facing leftward as during walking with head forward, i.e. the coherence peaked at contralateral heel strike at higher vertebral levels, whereas it peaked at ipsilateral heel strike at lower vertebral levels. As expected, the gains and coherences between EVS and EMG significantly decreased when walking with the head facing leftward.

The decrease in EVS-EMG coherence was asymmetric between the left and right sides of the trunk, i.e., the decrease was significant around left heel strikes at the left side between T9 and L4 and at the right side between T3 and T5. No effect of Head Orientation, Vertebral Level, Side of the trunk nor their interaction was found on the delays of the EMG responses. These results agree with the idea that less feedback control is required in the sagittal plane to stabilize walking. However, it does not explain the asymmetric nature of the decrease in coherence when walking with the head sideways.

### Decreases in coherence and gain may imply less feedback control in the sagittal plane

When standing with head facing rightward, EVS with the anode on the right mastoid process induces a backward sway as the response to an illusion of forward sway (Lund & Broberg, 1983; Britton *et al*., 1993; Ali *et al*., 2003; Mian & Day, 2014; Forbes *et al*., 2016). Underlying this backward sway, bilateral activation of posterior lower limbs and erectors spinae muscles was observed (Britton *et al*., 1993; Ali *et al*., 2003). Based on this, we expected a more symmetric spatiotemporal pattern of EVS-EMG coherence in paraspinal muscles during walking with the head facing sideways. In contrast, phasic EVS-EMG coherence with a significant decrease in magnitude was found when walking with the head facing to the left. Combined with the decrease in gains, the overall decrease indicates less reliance on vestibular signal for the control of the trunk muscles in the sagittal plane than in the frontal plane during walking.

This is likely be related to less reliance on feedback control in the sagittal plane, which is a consequence of the larger length than width of the base of support in walking and the fact that passively determined foot placement can stabilize gait in the sagittal, but not in the frontal plane (Bauby & Kuo, 2000; O’Connor & Kuo, 2009). When the head is turned, vestibular afferent signals partly encode movements in the sagittal plane, hence responses to EVS decrease compared to when facing forward. As reported in standing by Lund et al. (1983), with a head rotation of -60 degrees, EVS with the anode on the right, induced body sway in a direction of about -150 degrees. Thus, the remaining phasic EVS-EMG coupling pattern is possibly due to the projection of the disturbance signal on the frontal plane caused by the head rotation less than 90 degrees (Lund & Broberg, 1983; Britton *et al*., 1993; Ali *et al*., 2003).

### Asymmetry in the decrease in EMG-EVS coherence when walking while facing leftward

Asymmetric suppression in EVS-EMG coherence was observed between the left and right side of the trunk when walking with the head leftward. This is consistent with previous studies in lower limb muscles during standing (Britton *et al*., 1993; Ali *et al*., 2003; Dakin *et al*., 2007; Forbes *et al*., 2016). This asymmetric response was ascribed to the thorax rotation used to reach the large degree of head rotation without discomfort (Britton *et al*., 1993). Indeed, an averaged 5-degrees thorax rotation also exists in our study (HL-EVS: 5.191 ± 2.235 degrees, HL-No EVS: 4.886 ± 1.944 degrees, results not shown). This causes a biomechanical asymmetry, which possibly modifies the EVS induced activation of the paraspinal muscles.

Another potential explanation is that head turn induced an asymmetric projection of the vestibular signal in the reference frame of the trunk. The midpoint of the interaural axis between the vestibular labyrinths is assumed to form the origin of the vestibular reference frame in the transverse plane (Moore *et al*., 2005). However, the yaw axis of head rotation is located close to the atlanto-occipital and atlanto-axial joints, which are posterior to the vestibular labyrinths. Previously, in standing and sitting, the axis of yaw rotation was reported to be located about 10 millimetres posterior to the midpoint between the vestibular labyrinths (Moore *et al*., 2005).

With this distance, an offset of the vestibular axis to the left relative to the trunk reference frame will be induced (Figure 7). In this scheme, the vestibular signal encoding leftward motion would be reduced more than those encoding movement to the right, causing an asymmetric response to vestibular afference.

**Figure 7.**
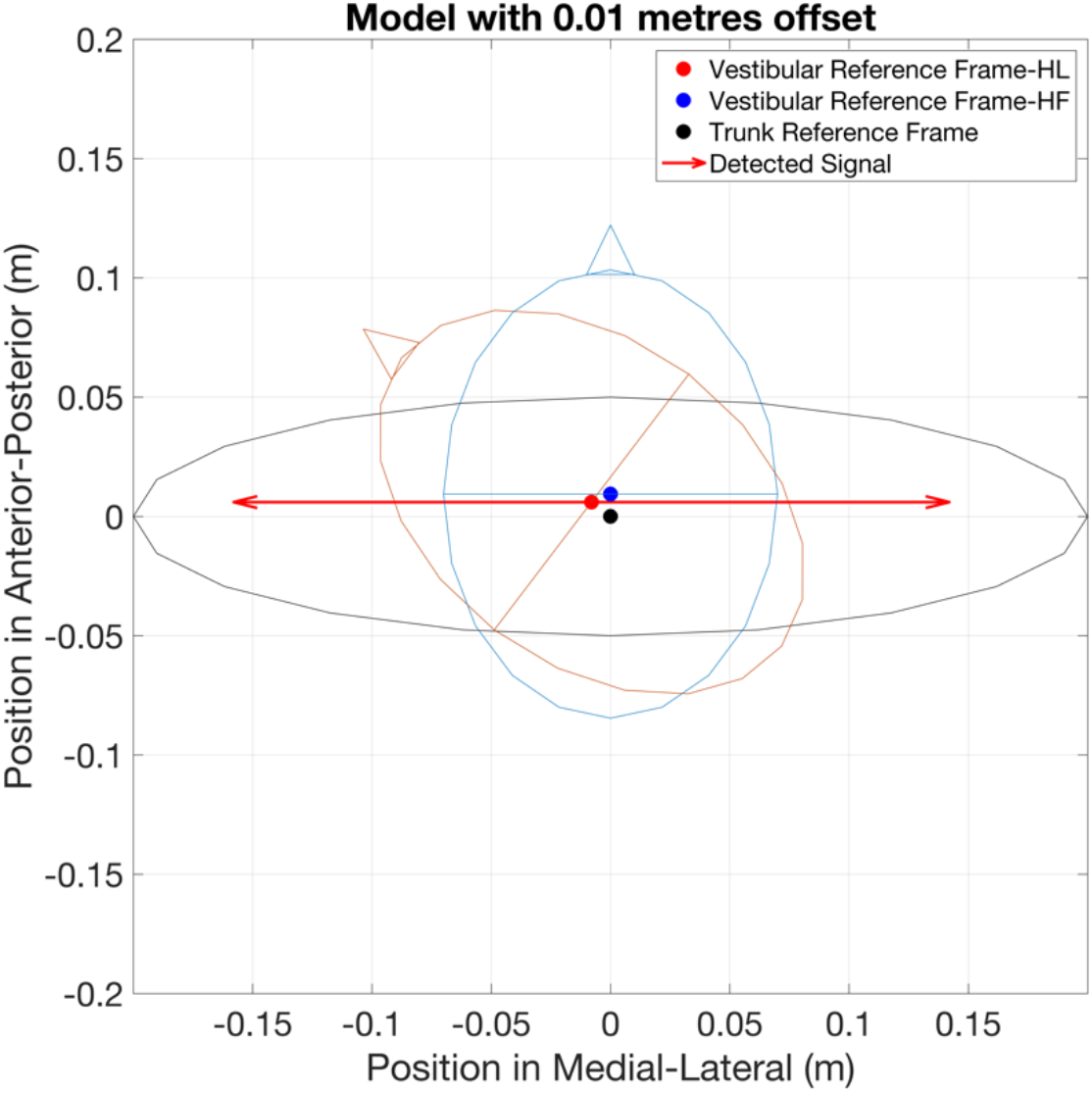
The head-fixed vestibular reference frame related to the orientation of the trunk (black circle) when facing forward (blue circle) and with the head turned 50 degrees (red circle). The X and Y-axis represent the position on the medial-lateral and anterior-posterior axes, in metres, respectively. Filled dots indicate the origin of trunk reference frame (in black), and of the vestibular reference frame when head facing forward (in blue) and facing leftward (in red). Vectors indicate the vestibular signals encoding illusions with the amplitude of 0.15 metres to both the left and right, detected in the vestibular reference frame with offset (Detected Signal, in red). With the offset, the projection of detected signals into the trunk reference frame induces an increased leftward illusion and a decreased rightward illusion. Consequently, compensatory responses to overcome the illusions are asymmetric in magnitude.

### Limitations

A limitation in this study is that the averaged head rotation did not reach 90 degrees, with which EVS would produce a disturbance purely in the sagittal plane (Forbes *et al*., 2016; Tisserand *et al*., 2018). However, as reported elsewhere, the maximal range of head yaw rotation is between ± 60° to ± 80° with slight thorax and/or pelvis rotation in standing (Ferrario *et al*., 2002; Tommasi *et al*., 2009; Higgins *et al*., 2023). The test conditions therefore reflect a feasible range of rotation. Additionally, only head leftward condition was included, which provides a limited view of the effect of head orientation on vestibular signal-based modulation of paraspinal muscle activity. Future investigations should include more orientations in both directions as was done for standing in Forbes et al (2016).

## Conclusion

When participants walked with their head facing leftward, significant phasic EVS-EMG coherence in paraspinal muscles was observed at ipsilateral and/or contralateral heel strikes, similar to when facing forward. Significant decreases in stimulation evoked responses (i.e., EVS-EMG coherence and gain) in paraspinal muscles were observed, when walking facing left compared to facing forward. This can possibly be explained by less need of the feedback control to stabilize walking in the sagittal plane compared to the frontal plane. The decrease in coherence was only significant around left heel strikes at the left side between T9 and L4 and at the right side between T3 and T5. The directionally specific response is likely be to a consequence of the offset between the origin of vestibular reference frame to head yaw rotation pivot. In sum, our findings confirmed the contribution of vestibular signal in trunk stabilization, while highlighting that this contribution is impacted by head orientation.

## Additional information

### Data Availability Statement

The original data are available from the corresponding author upon reasonable request.

### Author Contributions

Experiments were performed at the Dual Belt Lab in the Vrije Universiteit Amsterdam. Conception and design: Y.C.L., K.K.L., S.M.B., and J.H.D. Data acquisition: Y.C.L. Analysis and interpretation: Y.C.L., K.K.L., S.M.B., S.B., and J.H.D. Drafting and revising manuscript: Y.C.L., K.K.L., S.M.B., S.B., and J.H.D. All authors have read and approved the final version of this manuscript and agree to be accountable for all aspects of the work in ensuring that questions related to the accuracy or integrity of any part of the work are appropriately investigated and resolved. All persons designated as authors qualify for authorship, and all those who qualify for authorship are listed.

## Acknowledgements

The authors gratefully acknowledge Dr Patrick A. Forbes, Dr Mohammadreza Mahaki, Bert Clairbois and Richard Casius for their help. The authors also appreciate all participants for their voluntary participation.

